# Deep RNA-seq of male and female murine sensory neuron subtypes after nerve injury

**DOI:** 10.1101/2022.11.21.516781

**Authors:** Allison M Barry, Na Zhao, Xun Yang, David L Bennett, Georgios Baskozos

**Affiliations:** Nuffield Department of Clinical Neurosciences, University of Oxford, OX3 9DU, UK

## Abstract

Dorsal root ganglia (DRG) neurons have been well described for their role in driving both acute and pain. Although nerve injury is known to cause transcriptional dysregulation, how this differs across neuronal subtypes and the impact of sex is unclear. Here, we study the deep transcriptional profiles of multiple murine DRG populations in early and late pain states while considering sex. We have exploited currently available transgenics to label numerous subpopulations for fluorescent activated cell sorting (FACS) and subsequent transcriptomic analysis. Using bulk tissue samples, we are able to circumvent the issues of low transcript coverage and drop-outs seen with single cell datasets. This increases our power to detect novel and even subtle changes in gene expression within neuronal subtypes and discuss sexual dimorphism at the neuronal subtype level. We have curated this resource into an accessible database for other researchers (https://livedataoxford.shinyapps.io/drg-directory/). We see both stereotyped and unique subtype signatures in injured states after nerve injury at both an early and late timepoint. While all populations contribute to a general injury signature, subtype enrichment changes can also be seen. Within populations, there is not a strong intersection of sex and injury, but previously unknown sex differences in naïve states-particularly in A*β*-RA + A*δ*-LTMRs - still contribute to differences in injured neurons.

## Introduction

Chronic pain conditions affect up to 25% of the global population. Neuropathic pain, as a subclass, affects about 8%. It is directly tied to nervous system damage though trauma, disease, or therapeutic use (eg. chemotherapy, antiretrovirals) (***Costigan et al. (2009)***; ***Colloca et al. (2017)***). Current treatment options are widely considered inadequate and quality of life scores remain significantly reduced (***Attal et al. (2011)***; ***O’Connor (2009)***; ***James et al. (2018)***). Increasing lifespans, diabetic prevalence, and decreases in cancer mortality are all contributing to increases in these disorders, adding weight and urgency to deepen our understanding (***Colloca et al. (2017)***; ***James et al. (2018)***).

Primary afferent pathophysiology is thought to be a key driver for peripheral neuropathic pain disorders, with dorsal root ganglia (DRG) neurons being well described for their role in driving both acute and chronic pain. These neurons encompass a diverse collection of subtypes that are grouped by various factors, each intrinsically related. These include size, myelination, conduction velocity, projection patterns, end organ innervation, and functional properties (***Dubin and Patapoutian (2010)***; ***Handler and Ginty (2021)***).

More recently, single cell and single nuclear RNA-seq in mice (***Usoskin et al. (2015)***; ***LI et al. (2016)***; ***Zeisel et al. (2018)***; ***Nguyen et al. (2019)***) and human (***Nguyen et al. (2021)***; ***Tavares-Ferreira et al. (2022)***) have emphasized the diversity within DRG ganglia. Gene expression differs between these broad neuronal subpopulations and is indeed predictive of their functional properties (***Zheng et al. (2019)***). The diversity across subtypes is lost during bulk RNA-seq, due to the consolidation of all subtypes together. In single cell datasets, pseudo-bulk samples can be generated for each cluster, but this relies on a well-defined clustering that can be lost after nerve injury (***Hu et al. (2016)***; ***Nguyen et al. (2019)***). As such, changes at a subtype level remain unclear in painful states, and is the focus of the current study through the use of transgenic labelling and deep RNA-seq across populations.

Understanding sexual dimorphism in pain states is also a fundamental clinical issue. Females are much more likely to be living with chronic pain, and treatment efficacy can be sex-dependant (***Greenspan et al. (2007)***; ***Mogil (2012)***). In naïve states, quantitative sensory testing has also highlighted heightened pain sensitivity in females to a battery of acute noxious stimuli (***Bartley and Fillingim (2013)***). The case for studying sexual dimorphism is not new (***Berkley (1992)***; ***Unruh (1996)***), but historical biases within the research community resulted in a predominantly male focus (***Shansky (2019)***; ***Mogil (2012)***; ***Wald and Wu (2010)***).

More recent female-inclusive studies have revealed clear mechanistic differences between sexes at the immune and nervous system level. Sorge *et al*. report a prominent sex difference when studying mechanical hypersensitivity: While males depend on microglia activity, female animals depend on adaptive immune cells (***Sorge et al. (2011***, 2015)). Brain derived neurotrophic factor (BDNF) and prolactin both affect pain in a sex-dependant manner (***Patil et al. (2019)***; ***Moy et al. (2019)***) and there is evidence of higher level dimorphisms affecting pain percepts in the cortex (***Martin et al. (2019)***). At a transcript level, sexual dimorphism is also visible (***Baskozos et al. (2019)***; ***Mecklenburg et al. (2020)***), with transcriptomic differences seen in human DRG as well (***North et al. (2019)***; ***Tavares-Ferreira et al. (2022)***).

Here, we build on this research and address a gap through the deep transcriptional profiles of multiple murine DRG populations in acute and late pain states while considering sex differences. We have studied the molecular changes in five populations: *Scn10a*-expressing DRG, peptidergic and non-peptidergic nociceptors, as well as C-LTMRs and *Ntrk2*-expressing A-LTMRs. In a naïve state, we find subtype-specific sexual dimorphism in a small number of genes. This does not translate to a strong interaction of sex and injury, as the injury response seems to be consistent across sexes at the neuronal transcript level. We also see both stereotyped and unique subpopulation signatures in injured states after nerve injury at both an acute (3 day) and late (4 week) timepoint, with notable changes in C-LTMR and NP populations, as well as a distinct transcriptional program in A*β*-RA + A*δ*-LTMRs by 4 weeks compared to the other subtypes in the study. This data has been deposited in a searchable database at https://livedataoxford.shinyapps.io/drg-directory/.

## Results

Using transgenic labelling of neuronal DRG subtypes, 160 lumbar DRG samples were sequenced 3 days and 4 weeks after SNI (Fig 1A). This includes five neuronal subtypes sorted by fluorescence: general nociceptors, encoded by Scn10^cre^ (nociceptors), peptidergic nociceptors from Calca^creERT2^ (PEP/peptidergic), non-peptidergic nociceptors by Mrgprd^creERT2^ (NP/non-peptidergic), C-low threshold mechanoreceptors encoded by Th^creERT2^ (C-LTMRs) and Ntrk2^creERT2^ expressing LTMRs (Rapidly adapting (A*β*-RA) and D-Hairs (A*δ*-LTMRs)) (Fig 1B-C, Supplemental Figure 1). We recognize that our general nociceptor population expressing *Scn10a* does not exclusively comprise high threshold afferents. C-LTMRs are included within this subtype, based on the co-expression of *Th* and *Scn10a*. They make up a much smaller proportion of overall cells than the peptidergic and non-peptidergic nociceptor subpopulations, resulting in a “nociceptor-like” population.

**Figure 1.**
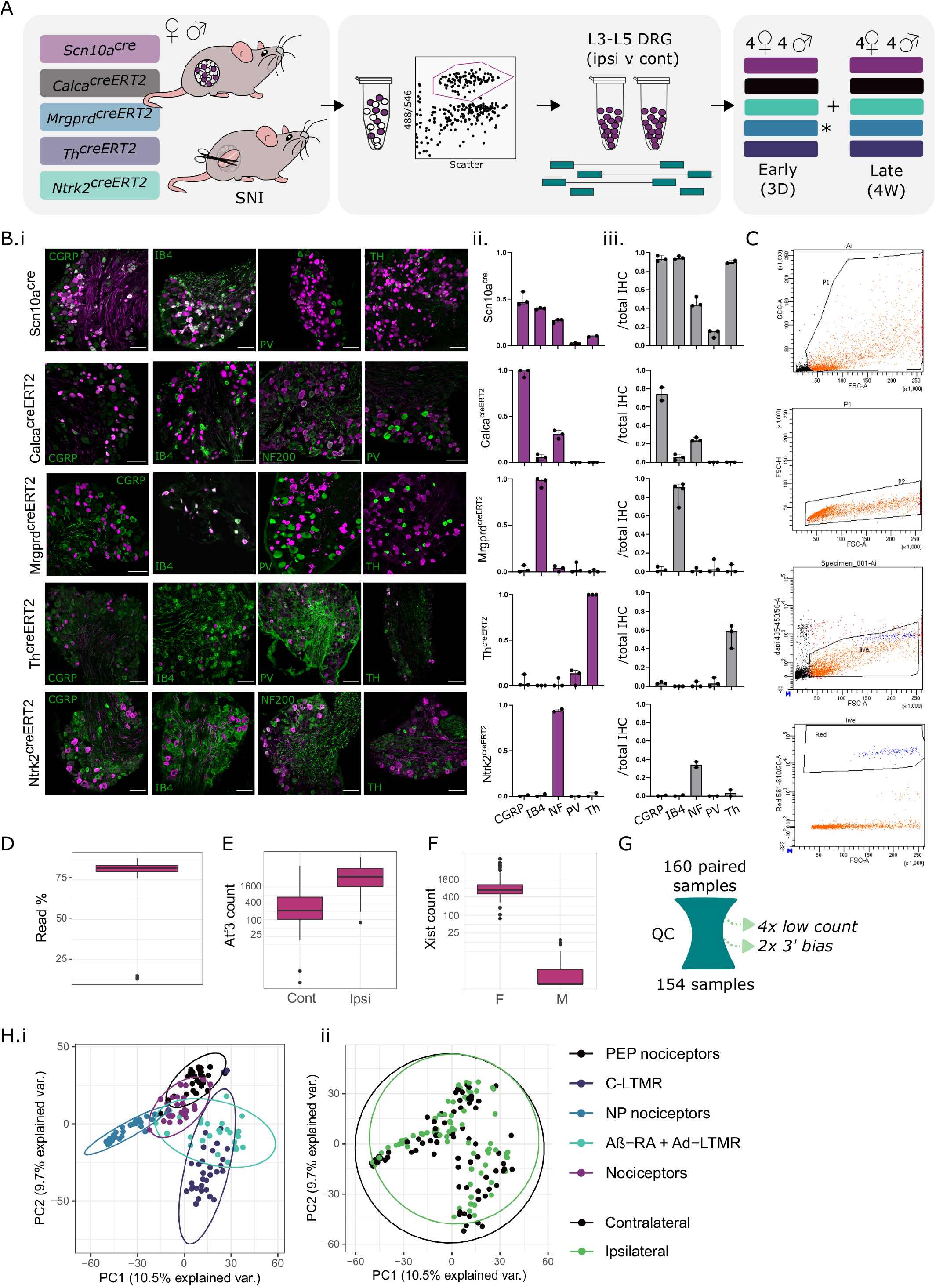
Experimental overview for mouse subtype RNA-seq of five neuronal subtypes after nerve injury. A. Overview schematic, highlighting five transgenic mouse lines used to label and sort “bulk” subtype samples for downstream sequencing. Males and females were collected 3 days (3D) and 4 weeks (4W) after spared nerve injury (SNI). B Transgenic validation of Scn10a^cre^, Calca^creERT2^, Mrgprd^creERT2^, Th^creERT2^, and Ntrk2^creERT2^ lines. B.i. example IHC, B.ii. IHC overlap with reporter line. B.iii Reporter overlap with IHC. C. Samples FACS gating (*Mrgprd*+ cells, gating for scatter, live/dead, and tdTomato), with addition details in 1. D. Percentage of uniquely mapped reads by sample. E. *Atf3* raw count data. F. *Xist* raw count data. G. Schematic of QC. 154 samples passed. H. PCA biplot by subtype (i) and injury status (ii). Plots uncorrected for batch are shown in 2. **Figure 1 —figure supplement 1**. Fluorescence activated cell sorting of sensory neuron subtypes **Figure 1 —figure supplement 2**. Supplemental sample clustering

Together, 154 samples passed QC, removing samples with low read counts or 3’ bias. Male and female samples are clearly distinguishable by sex-linked genes such as *Xist*, and ipsilateral (“injured”) samples can be distinguished from contralateral controls by key injury markers such as *Atf3* (Fig 1D-G). A batch effect was introduced on the first sample collection day, affecting paired (ipsilateral and contralateral) samples for Calca^creERT2^ and Scn10a^cre^ females (Supplemental Figure 2). We controlled for this effect in all downstream analyses (see methods).

Sensory neurons undergo broad, stereotyped changes after injury (***Nguyen et al. (2019)***). Even so, samples largely cluster by neuronal subtype across conditions (Fig 1H). While the bulk subpopulation methodology employed here allows deep sequencing within populations, each resulting sample contains a mix of injured and intact neurons from ipsilateral ganglia. This cell mixture likely dampens the stereotyped changes seen previously in single cell RNA-seq (***Nguyen et al. (2019)***).

Analyses were first performed on contralateral (“naïve”) samples. As expected, samples initially cluster by batch before clustering by subpopulation (Supplemental Figure 2C-D). General nociceptors (nociceptors), as well as peptidergic and non-peptidergic nociceptor subpopulations largely separate from C-LTMRs and *Ntrk2*-labelled A*β*-RA + A*δ*-LTMRs.

### Contralateral (“naïve”) samples match previous data

In line with previous reports, hallmark gene expression can be seen within each population (Figure 2, Supplemental Table 1). Voltage gated sodium (*Scn*) channels, transient receptor potential (*Trp*), Gamma-aminobutyric acid (*GABA*) receptors (*Gabra*), and two pore potassium channels (*Kcnk*) show varying subtype specificity (2B-C).

**Figure 2.**
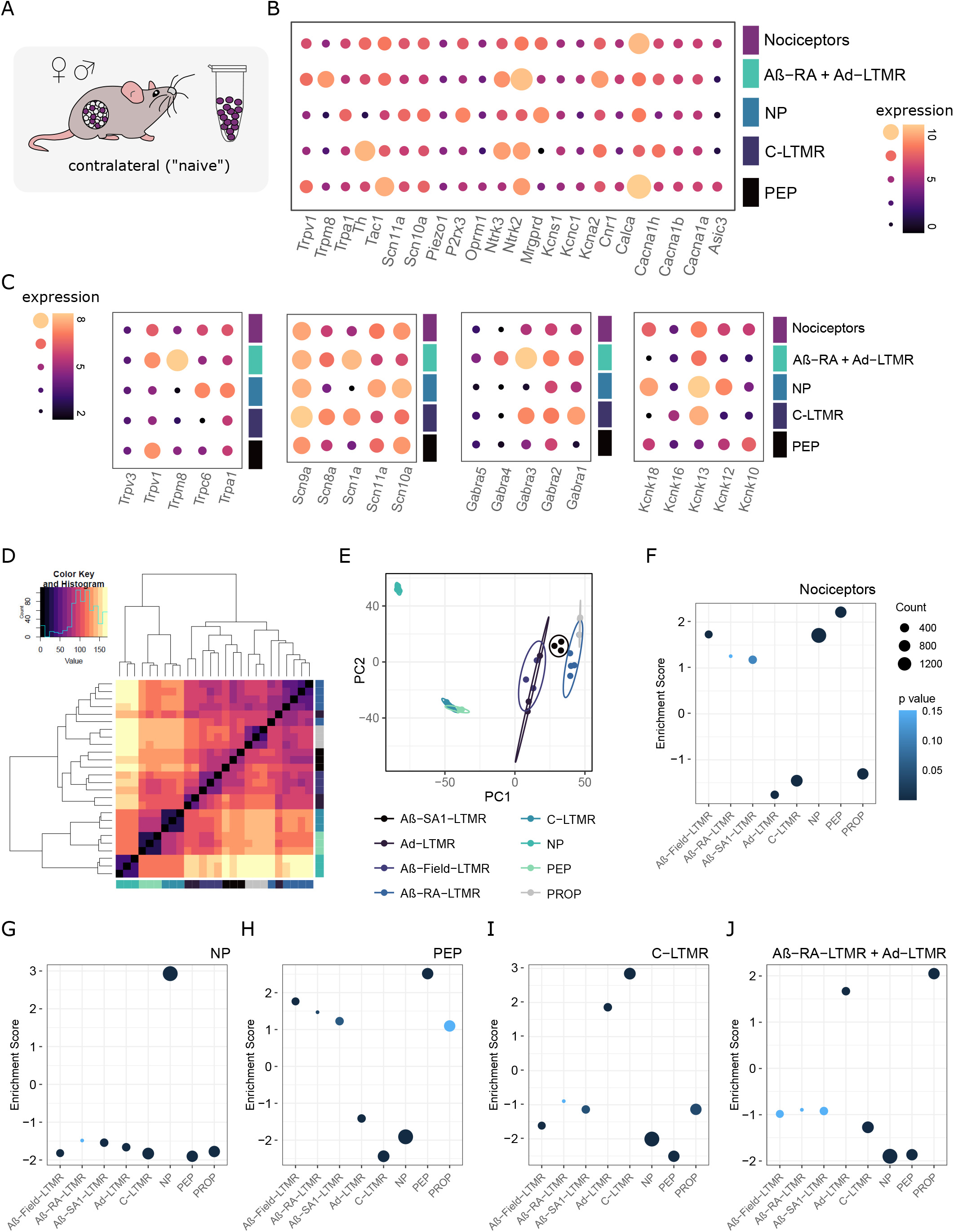
RNA-seq validation against previously published work, combining male + female samples. A. Contralateral tissue was compared to previously published naïve datasets. B. Hallmark gene expression across contralateral samples. Expression plotted as VST transformed count data.C. Ion channel expression across contralateral samples. D-E. Zheng *et al*. 2019 naïve subpopulation clustering (mixed sex). F-J: Subtype enrichment against gene sets derived from Zheng *et al*. 2019 (see methods for details, gene sets provided in Supplemental Table 3). F (nociceptors), G (peptidergic nociceptors), H (non-peptidergic nociceptors), I (C-LTMRS), and J (A*β*-RA + A*δ*-LTMRs). Plotted as normalized enrichment scores, coloured by p-value. Full lists of scores and p-values are available in Supplemental Table 2.

*Scn10a* and *Scn11a* are enriched in high threshold populations, whereas *Scn1a* is enriched in LTMRs. *Mrgprd* and *Mrgpr* family members, *P2rx3*, *Pirt*, and *Trpa1* are enriched in non-peptidergic nociceptors, in line with previous reports. *Ntrk2*, *Scn1a*, and *Trpc1* are enriched in A*β*-RA + A*δ*-LTMRs, whereas *Th*, *Tafa4*, *Gfra2*, and *Slc17a8* (VGLUT3) are all enriched in C-LTMRs. Key peptidergic markers such as *Calca* and *Trpv1* do not hit our enrichment filtering criteria, likely due to strong expression in both peptidergic and general nociceptor populations.

Two-pore potassium channels (K2Ps) have been implicated in various pain conditions, including the role of *Kcnk18* (TRESK) in migraine (***Royal et al. (2019)***; ***Li and Toyoda (2015)***). We see a range of subpopulation enrichments for this gene family in our dataset. *Kcnk13* (THIK-1), *Kcnk12* (THIK-2), and *Kcnk18* (TRESK) are all enriched in non-peptidergic nociceptors. *Kcnk16* (TALK-1) is enriched in C-LTMRs. *Kcnk10*, encoding the TREK-2, has previously been implicated in spontaneous pain in rats, with expression limited to IB4-binding neurons (***Acosta et al. (2014)***). Here, we see higher expression in peptidergic nociceptors. This peptidergic-enrichment profile is supported by the transcriptional data published by Zheng and colleagues (***Zheng et al.(2019)***).

To validate our sequencing approach, gene enrichment analyses were performed against previously published naïve subtypes (Figure 2D-J, Supplemental Tables 2-3). Count data were log transformed using DESeq2, mirroring our analysis to generate eight subpopulation-specific groups, as defined by Zheng and colleagues (***Zheng et al. (2019)***).

Our samples correlate strongly to this data. Our *Scn10a* population appears to be largely nociceptor, with positive enrichment for both PEP and NP populations (Fig 2F). Negative enrichment is seen for C-LTMRs, likely reflecting subpopulation proportions within this broad grouping. Our NP samples are positively enriched for the NP gene set, and negatively enriched for all other gene signatures (Fig 2G). To complement, our PEP samples show strong positive enrichment for PEP, as well as negative enrichment for NP and C-LTMR signatures (Fig 2H). Unlike our peptidergic population, our general nociceptor is negatively enriched for the proprioceptive signature. Both our nociceptor and peptidergic nociceptor populations show negative enrichment for the AA*δ*-LTMR gene set, but positive enrichment of A*β*-Field-LTMR signatures.

C-LTMRs are highly enriched for the C-LTMR gene set, along with a positive enrichment of A*δ*-LTMRs (Fig 2I). Correspondingly, this population shows negative enrichment of high threshold populations (NP and PEP), along with the proprioceptive and A*β*-Field-LTMR signatures.

A*β*-RA + A*δ*-LTMRs show positive enrichment for the proprioceptive gene set, followed by enrichment for A*δ*-LTMRs (2J). As expected, this population is negatively enriched for high threshold signatures. Enrichment scores and adjusted p-values are listed in Supplemental Table 2.

Together, these enrichments lend confidence to our methodology and support the use of this dataset to interrogate population sex differences and as a baseline against injured neurons.

#### Naïve sex differences across subtypes

With many clinically-relevant pain conditions showing sexual dimorphisms, there is a keen interest to explore sex differences within each DRG subtype transcriptome. Across subpopulations, most genes are expressed to similar levels in males and females (Fig 3). DEGs are defined here as an FDR < 0.05 and an absolute log_2_ fold change (LFC) > 1. From this, only six genes were significantly regulated between males and females in all populations, each X- or Y-linked (*Kdm5d*, *Uty*, *Ddx3y*, *Eif2s3y*, *Tsix*, *Xist*). *Gm29650*, which is also sex-linked, was differentially expressed in 4/5 populations. The only other DEG shared across any subtypes is *Sprr1a*, which is more highly expressed in male nociceptors and non-peptidergic nociceptors, and may reflect differences in naïve cells, or minor wounds from in-cage fighting which occur at higher rates in males.

**Figure 3.**
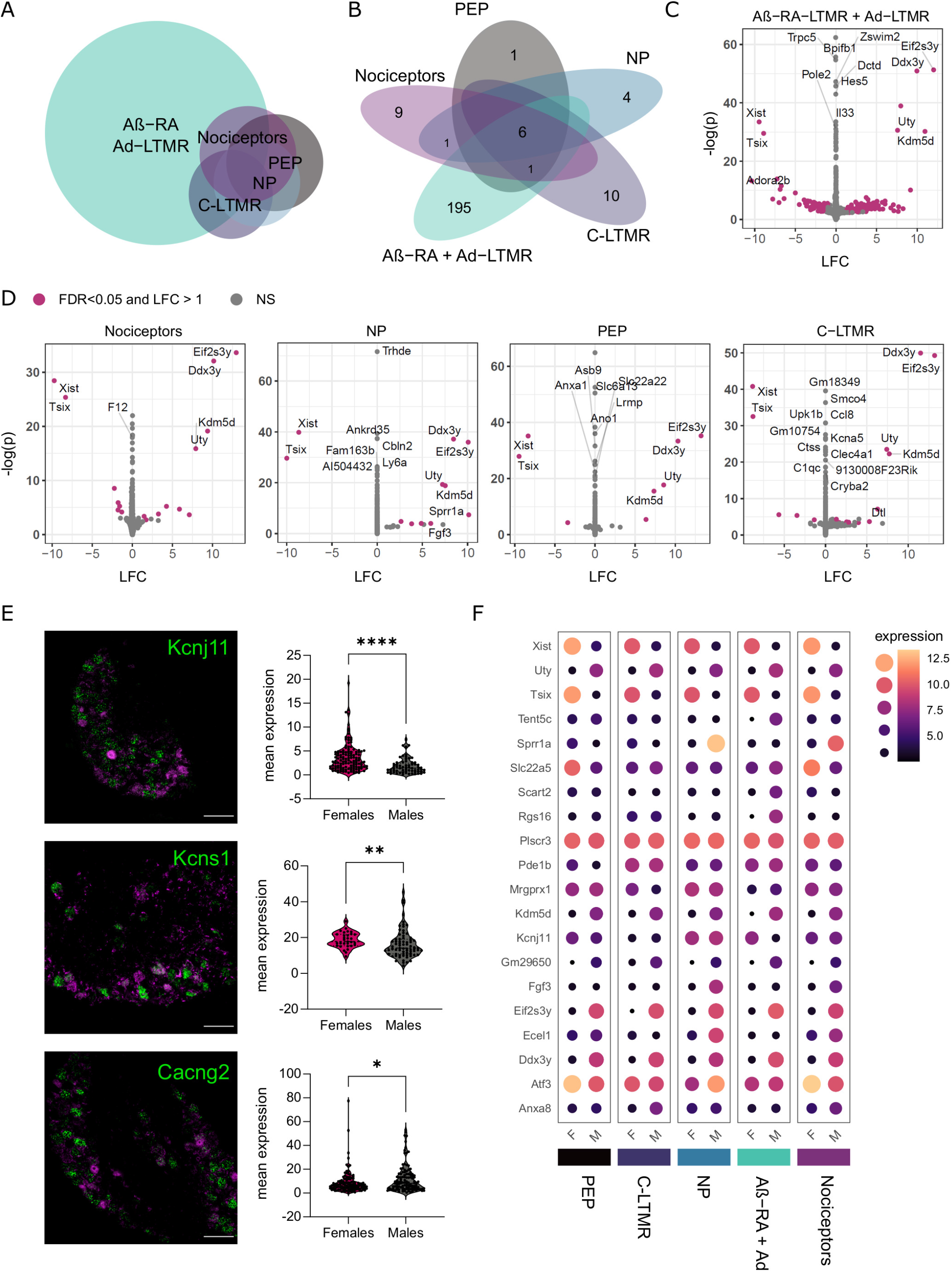
Few sex differences are seen in uninjured neuronal subtypes, with the majority in A*β*-RA + A*δ*-LTMRs. Few sex differences are seen in uninjured neuronal subtypes, with the majority in *Ntrk2*+ LTMRs. A. Euler plot for sexually dimorphic genes. B. Number of sexually dimorphic genes within each subpopulation examined (FDR < 0.05, LFC > 1). C-D. Contralateral samples show differential gene expression across sexes (male vs female, ie. LFC > 0 = upregulated in males). E. *In situ* validation of gene candidates regulated in A*β*-RA + A*δ*-LTMRs (n=3 mice, Mann Whitney test. Data points represent individual cells). Top to bottom: *Kcnj11* (p < 0.0001), *Kcns1* (p = 0.0054), *Cacng2* (p = 0.0242) (green) with *Ntrk2* (magenta). Note. *Cacng2* RNA-seq suggests upregulation in females (opposite). F. Dot plots highlighting key DEGs, plotted as median transformed counts.

This absence of a large sensory-neuron wide sex signature is consistent with previous work in mice, where all but one DEG reported in adult, lumbar DRG were X- or Y-linked (***Smith-Anttila et al. (2020)***). Nine autosomal genes were found to be regulated in sacral DRG (***Smith-Anttila et al. (2020)***), and these do not overlap with the DEGs reported within populations here, with the exception of *Clvs1* in A*β*-RA + A*δ*-LTMRs.

At a subpopulation-level, a stronger sexual dimorphism emerges. Across populations, many genes hit an FDR < 0.05, but moderated fold changes suggest a negligible effect (near 0 LFC) in these genes (Supplemental Table 4-5). The majority of DEGs are seen in lowly expressed genes within unique neuronal subtypes. We see predominant transcriptional changes within A*β*-RA + A*δ*-LTMRs (202 genes with FDR < 0.05, LFC > 1), with few differences in the other populations. Here, RNA-seq and *in situ* hybridization validation of *Kcnj11* and *Kcns1* (Fig 3E) suggest they are more strongly expressed in female TRKB+ LTMRs. *KCNS1* has previously been suggested as a marker of LTMRs in humans (***Tavares-Ferreira et al. (2022)***), and cross-species validation of possible sexual dimorphism in TRKB+ and other LTMR populations is recommended. Not all candidates were validated using *in situ* via naïve, wildtype controls. Of the three candidates explored, *Cacng2* (Stargazin) is significantly upregulated in males via *in situ* (3E) but downregulated in our RNA-seq results (Supplemental Table 4). This still suggests sexual dimorphism within this LTMR population, but warrants note as it may be a result of population labelling via *in situ* vs transgenics or a difference in contralateral vs naïve tissue.

GO term analyses highlight enrichment for ion channel transport and transmembrane transport in females, although few genes are implicated in each (Supplemental Table 6). GSEA analyses against “all gene sets” available from Molecular Signatures Database (MSigDB) show no enrichments in this population (Supplemental Table 7).

In the four other populations studied, most GO terms centre around sex-linked processes. Other relevant GO pathways include the detection of temperature stimulus involved in sensory perception of pain (nociceptors), immune response and cholinergic synaptic transmission (PEP), regulation of sensory perception of pain (NP), and chemosensory behavior (C-LTMR). All supplemental tables for DEGs and pathway analyses, including full DEG tables, GO, and GSEA analyses for each subpopulations are available in the supplementals.

#### General injury signatures

General injury signatures were examined by combining samples across subtypes (Fig 4, Supplemental Table 8-9). These samples will be biased toward nociceptors, due to the inclusion of three nociceptor populations (Scn10a^cre^, Calca^creERT2^, and Mrgprd^creERT2^) and two LTMR populations (Th^creERT2^ and Ntrk2^creERT2^) in equal numbers.

**Figure 4.**
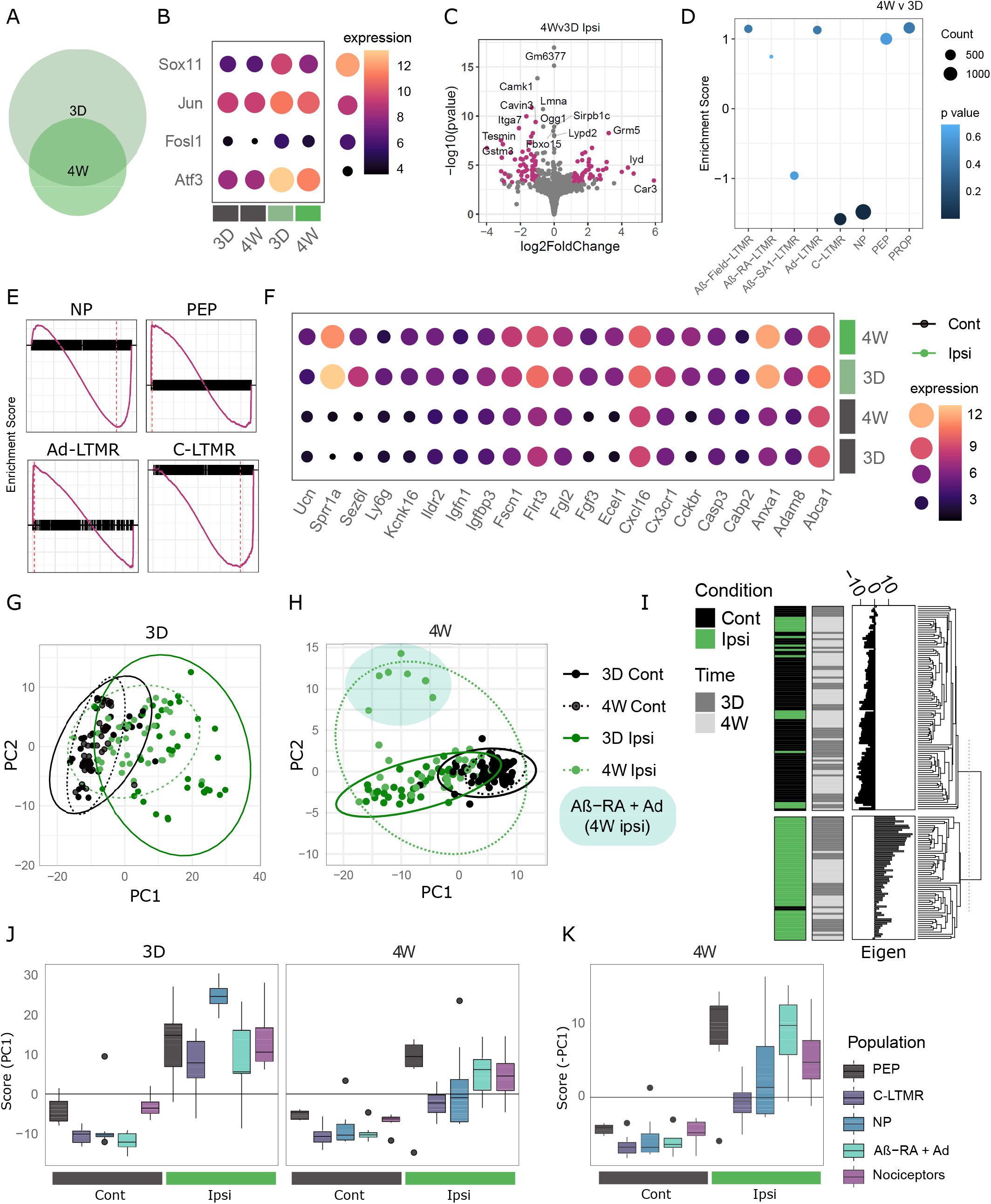
General injury mapping shows stereotyped changes and subtype differences across timepoints. A. Euler plot showing differentially expressed genes after SNI at 3 days (3D) and 4 weeks (4W) for combined subtypes. B. Key injury markers are upregulated across ipsilateral samples at both timepoints, plotted as median VST expression across groups. C. Injured samples (ipsi) show differential gene expression across timepoints (4W v 3D). D-E. GSEA analysis of subtype enrichment between 4W and 3D ipsilateral samples. NP and C-LTMR signatures are significantly reduced at 4 weeks, with values listed in Supplemental Table 12. E. Enrichment plots for key subtypes, by enrichment score for ranked genes (black). F. Example differentially expressed genes (DEGs) shared across timepoints, plotted as median VST expression across groups. G-H. Supervised PCA bi-plot for DEGs at 3 days (G) and 4 weeks (H). I. Dendrogram split by k-means of 2, highlighting positive and negative injury scores from 3D DEGs largely correlate to sample condition. J-K. PC1 correlation across subtypes and time. Boxplot whiskers show 1.5 IQR. J. 3D signature: All five subtypes shows a significant difference between ipsilateral and contralateral samples at both timepoints (Kruskal-Wallis Rank Sum, followed by pairwise Wilcoxon with BH correction against a grouping factor (population, condition, and timepoint)). Only NP and C-LTMR ipsilateral samples are different between 4 weeks and 3 days (FDR = 0.00185 and 0.00995 respectively), reflecting their return towards baseline. K. 4W signature: all five subtypes show a significant difference between ipsilateral and contralateral samples, plotted as -score for visualization with J.

At both 3 days and 4 weeks, we see predominant upregulation of genes associated with classical injury signatures, including *Atf3*, *Jun*, *Sox11*, and *Fosl1* (4B, Supplemental Table 8). The overall number of DEGs (LFC > 1, FDR < 0.05) is reduced over time, from 521 at three days to 162 by four weeks. Of these, 96 are shared across timepoints (Supplemental Table 9), with some highlighted in Figure 4C. Subtype-enriched genes appear acutely downregulated at three days, in line with previous reports although few genes reach our significance threshold. This is likely due to the high variability from combining samples across populations.

Our timecourse was selected to highlight the progression from a more acute to a later injury state after SNI (Fig 4C-D). We can probe this transition by comparing ipsilateral samples across timepoints. At four weeks, we see the downregulation of *Atf3*, as well as an upregulation of some subtype-specific genes towards baseline, such as *Ntrk2*. *Slc17a7* (VGLUT1), typically a marker of larger diameter DRG is positively enriched at 4 weeks compared to 3 days after SNI (LFC = 2.25, p.adj = 1.7E-07), although this change is not reflected by a median expression change and is likely driven by a subgroup of samples. For other genes, such as *Npy*, gene expression increases from 3 days to 4 weeks, where it is significantly higher than contralateral levels, suggesting a more long term change.

In a more acute state, GO analyses show enrichment in the regulation of cell population proliferation, positive regulation of apoptotic process, and inflammatory response after injury (Supplemental Table 10). Many of these processes remain enriched at 4 weeks, even with the overall reduction in DEGs. GSEA enrichment at 3 days also shows downregulation of electron transport and oxidative phosphorylation paired to a positive enrichment of inflammation, receptor regulator activity, and cell migration (Supplemental Table 11). When comparing ipsilateral samples over time, GO analyses also suggest functional changes. Three day injured samples show enrichment of apoptotic process, cytokine response and, and positive regulation of gene expression. In a later state, there is an enrichment for protein import, long-term memory, and the regulation of long-term neuronal synaptic plasticity. Taken together, these results suggest we are accurately capturing injury signatures across our dataset.

#### Injury phenotypes by subtype

A major strength of this study is the ability to probe subtype specific patterns in a murine model of neuropathic pain. Injured neurons were previously shown to lose cell-type specific identities after nerve injury in a time-dependant process (***Renthal et al.*** (2020)). At 4 weeks post-SNI, injured samples show a negative enrichment for C-LTMRs and NP nociceptors compared to their 3 day injured counterparts (Fig 4D-E). Here, we are also able to explore subpopulation-specific and common injury signatures across cell types (Fig 4G-K), before contrasting samples within each population (Fig 5).

**Figure 5.**
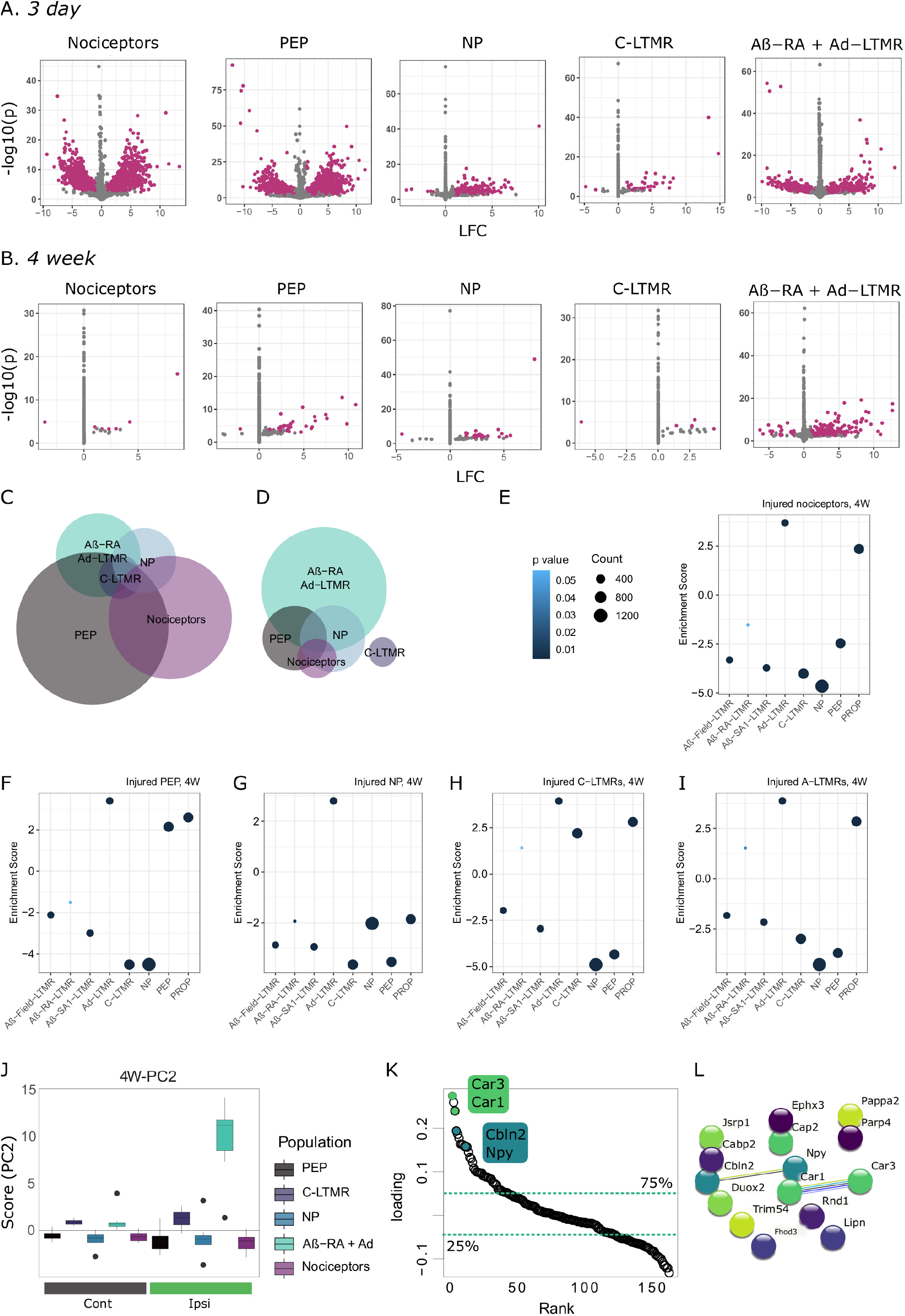
Subtype injury mapping shows stereotyped changes and subtype differences across timepoints. Subtype injury mapping shows stereotyped changes and subtype differences across timepoints. A-D: Differential gene expression across timepoints and subtypes. Volcano plots for 3 day (A) and 4 week (B) subtypes, with DEGs highlighted in magenta. Euler plots showing DEG overlap at 3D (C) and 4W (D). E-I: GSEA analyses against previously published data (***Zheng et al. (2019)***) reveals a lack of clear subpopulation signatures in injured nociceptors and NP nociceptors by 4 weeks (scores listed in Supplemental Table 18). All subtypes show a significant enrichment for A*δ*-LTMRs, which was not seen in naïve nociceptor populations. Naïve data is shown in Fig 2. J-L: Subtype-specific signatures extracted from 4W SPCA presented in Figure 4 shows specificity for Ntrk2-injured neurons. J. PC2 correlation across subtypes four weeks after SNI. Boxplot whiskers show 1.5 IQR. K. Ranked loadings (PC2) for all DEGs at 4 weeks after injury. Dashed lines highlight quartiles. L. STRING database interactions for top 15 DEGs (ranked by loadings).

To start, an acute injury signature was extracted from our initial list of DEGs at 3 days (from combined samples, above) through a supervised PCA (SPCA) and compared across subtypes (Fig 4G,I-J). This provides an unbiased signature through the linear combination of individual gene expressions. All five subtypes studied show a significant difference (FDR < 0.05) between ipsilateral and contralateral samples at both timepoints (Kruskal-Wallis Rank Sum, followed by pairwise Wilcoxon with BH correction against a grouping factor (population, condition, and timepoint)). Only NP and C-LTMR show a significant difference between 4 weeks and 3 days (FDR = 0.00185 and 0.00995 respectively), reflecting their stronger return towards baseline.

Eigenvectors were also extracted from an SPCA on 4W DEGs to form a late injury signature across populations (Fig 4H). This signature is driven largely by general nociceptors, peptidergic nociceptors, and A*β*-RA + A*δ*-LTMRs, although all subtypes show a significant difference from their contralateral counterparts at 4 weeks (Kruskal-Wallis Rank Sum, followed by pairwise Wilcoxon with BH correction) (Fig 4K).

#### Differential gene expression changes by subtype

Within population testing for differentially expressed genes show a number of shared regulated genes typical of injury signatures (including *Atf3*, *Sprr1a*, and *Sox11*), as well as a subset of subtype-specific DEGs (Fig 5A-D, Supplemental Table 13-18). Over 5000 genes are regulated overall and are mostly driven by changes in the general nociceptors (2620) and peptidergic nociceptor (3270) samples at three days. Fewer DEGs are seen in other populations, with 179 DEGs for non-peptidergic nociceptors, 640 DEGs for A*β*-RA + A*δ*-LTMRs, and only 36 DEGs in C-LTMRs (with an LFC > 1, FDR < 0.05). Using an LFC cutoff of 1, we are focusing on genes with good expression changes thought to correspond to biological relevance. Even so, small changes in gene expression can be biologically relevant, and full results tables are provided online. With the populations studied, we do not see a clear LTMR-specific pattern: no genes exclusively regulated in both C-LTMRs and A*β*-RA + A*δ*-LTMRs.

Most genes are either regulated in the same direction, or only regulated in a subset of populations. Even so, over 400 DEGs are regulated in opposing directions, which may provide a unique look at subtype differences after injury (Supplemental Table 14). For example, *Ints5* is an integrator complex involved in RNA transcription which is upregulated in PEP and general nociceptors while being downregulated in A*β*-RA + A*δ*-LTMRs. The GTP binding protein *Gtpbp1* has been previously implicated in neuronal death through translational regulation (***Terrey et al. (2020)***). Here, we see subtype differences in regulation, being upregulated in PEP and A*β*-RA + A*δ*-LTMRs, but downregulated in general nociceptors (with downward trends in NP and C-LTMRs). Other genes showing bidirectional regulation include *Rnd1*, or Rho Family GTPase 1, which has been discussed previously for its role in axon outgrowth, while *Wnk4* is related to actin cytoskeleton remodelling by Rho GTPases, as well as ion channel regulation.

At four weeks post-SNI, there is a reduced number of DEGs across all subtypes, compared to our 3 day results (Fig 5B). The most changes are present in A*β*-RA + A*δ*-LTMRs (217 DEGs). Non-peptidergic and peptidergic nociceptors show the next highest number of DEGs, with 25 and 36 respectively (sharing *Cckbr*, *Tubb6*, *Atf3*, *Gpr151*, *Wt1*, *Pde6b*, *Cyp26a1*, and *Cdk6*). Few changes are seen in our general nociceptor population, with only 7 DEGs with a moderated LFC > 1 (*Cckbr*, *Ttll10*, *Phox2b*, *Rpl31-ps13*, *S100a8*, *Gata5os*, and *Gm47138*). C-LTMRs also show few changes (5 DEGs), in line with the acute signature (*Hbaa1*, *Nefh*, *S100b*, *Adtrp*, *Gm35097*). No genes show regulation in opposing directions across subtypes by 4 weeks, although some, like *Rnd1* remain regulated.

Many studies highlight cell-type or injury-specific gene regulation mechanisms, as they regulate important mechanisms of neuropathic pain. Additionally, they provide possible new avenues to target and modulate neurons in injured states. Our current dataset is well suited for these enquires. Across timepoints, a number of DEGs correspond to transcription factors (“GO:0003700”) and/or are involved in gene regulation (“GO:0010468”). This includes shared regulators like *Atf3*, *Hoxa2*, *Twist2*, *Cdk6*, *Prdm10*, and *Trim34b* as well as numerous subtype-specific regulators (Supplemental Tables 16-17).

Next, we queried cell type enrichment in our injured samples (Figure 5E-I, Supplemental Table 18). By 4 weeks post-SNI, GSEA enrichment of subpopulation signatures varies from their naïve counterparts, previously discussed in Fig 2. Injured nociceptors and NP nociceptors no longer show clear subpopulation delineations, while injured PEP and C-LTMRs both show enrichment for their respective populations. Across all subtypes, there is a positive enrichment in the A*δ*-LTMR signature, previously only seen in naïve LTMR samples. Injured A*β*-RA + A*δ*-LTMRs also show a new negative enrichment for A*β*-Field and A*β*-SA1-LTMRs.

Using a general injury signature in Fig 4, a subset of samples shared variation in PC2. This eigenvector was also extracted for comparison (Fig 5J-L). Here, this variation is driven by A*β*-RA + A*δ*-LTMRs ipsilateral samples, suggesting a distinct injury signature in this population not captured in the other subtypes. Gene loadings for PC2 were extracted and ranked by loading for further analyses (5K), with the top and bottom 25% quartiles extracted for GO analyses. Both “response to stimulus” and “actin filament organization” were upregulated GO terms, while “immune response” and “synapse maturation” were downregulated. No GSEA enrichments on ranked loadings were present.

The STRING database was next queried for possible interactions between gene products. In the top quartile (41 DEGs), interactions are seen between *Car1-Car3*, *Unc-Npy-Cbln2*, *Hrk-Pmaip1*, and *Uox-Mbl2*. Interactions for the genes with the highest loadings are highlighted in Fig 5L.

#### Sexual dimorphism in injured states

To see if male and female sensory neurons differ in their maintenance of later neuropathic pain states, we fitted an interaction model for sex and condition (Fig 6). Acute effects were not studied due to the colinearity of the batch and sex in a subset of the populations. Using this stringent modelling, which requires genes to be regulated in injury, as well as have a differential response to sex, we detected no differences when pooling populations at 4 weeks, or when subsetting our data to interrogate within subtype (Fig 6A). The number of genes with FDR < 0.05 range from 9 (NP) to 212 (A*β*-RA + A*δ*-LTMRs), but moderated fold changes centre towards zero, suggesting these result from the high variability in low count genes, instead of biologically meaningful differences.

**Figure 6.**
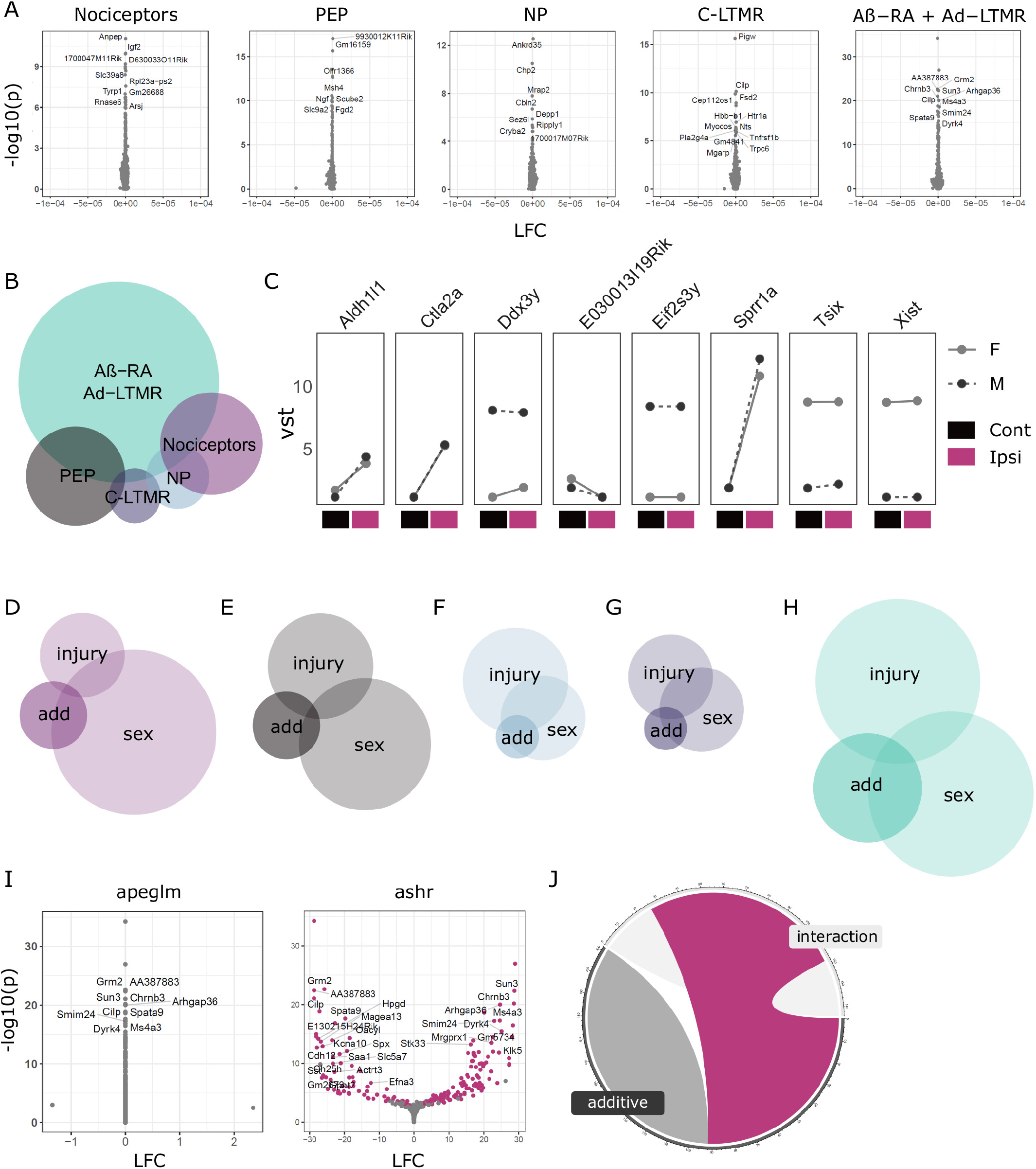
Sexual dimorphism in neuronal subtype injury responses. A. Transcriptomic analyses in primary afferents reveal no clear interaction of sex and injury 4 weeks after SNI. B. Euler plot of DEGs using an additive model contrasting sex and injury differences. C. Line plots of DEGs shared across at least two subtypes. D-H: Across subtypes, DEGs from this additive modelling (“add”) appear to be driven partly by sex differences in basal expression levels (“sex”), as well as some overlap with genes generally regulated in injury (“injury”). D: General nociceptors. E: PEP nociceptors. F: NP nociceptors. G: C-LTMRs. H. A*β*-RA + A*δ*-LTMRs. I. Example volcano plots for the interaction of sex and injury for A*β*-RA + A*δ*-LTMRs, with *apeglm* and *ashr* shrinkage. J. 42% of regulated genes are shared across our interaction and additive models (magenta) which are not regulated with *apeglm*.

By definition, interaction effects are calculated using the reference mean expression from only one sex. This can mask effects in genes which are lowly expressed in one sex during control states. To overcome this, we further explored sexual dimorphism using a more relaxed additive design, subtracting sex and injury comparisons to pull out sex differences which cannot be explained by the injury effect alone (Fig 6B-J, Supplemental Table 19).

Using this approach a number of DEGs are seen, with the majority in A*β*-RA + A*δ*-LTMRs (144 genes), followed by peptidergic nociceptors (41) and general nociceptor (34) populations (Fig 6B).

Eight genes show regulation in multiple subtypes. This includes a subgroup of sex-linked genes (eg. *Xist*, *Tsix*, *Kdm5d*), as well as those regulated in injured states, like *Sprr1a*. Only one, *Tsix*, shows regulation across all subtypes, but this does not appear to be a strong interaction of sex and injury (Fig 6C). Across subtypes, DEGs from this differential response to injury appear to be driven partly by sex differences in basal expression levels, as well as some overlap with genes generally regulated in injury.

A*β*-RA + A*δ*-LTMRs show the largest number of DEGs. Here, there is a 32% overlap with significantly regulated genes from naïve male vs female samples and a 12% overlap with genes regulated in an injured state (6H). Together, 40% of DEGs overlap with significantly regulated genes for sex in control samples or injury at 4 weeks, based on an FDR < 0.05. This does not account for fold changes, due to differences in shrinkage methods (*apeglm* vs *ashr*, see methods).

To see if the difference between our interactive and additive modeling was primarily a caveat of shrinkage priors, our interaction fold changes were re-calculated using *ashr*, where 42% of DEGs detected through the additive model overlap (Fig 6I-J). Shrinkage priors aim to limit noise within a dataset, and plotting non-transformed counts for these genes highlights the variability present. Some fold changes appear driven by a single sample per condition, while others show stronger trends. This highlights how the variability in lowly expressed genes can limit conclusive analyses.

Taken together, we interpret this as a lack of strong sexual dimorphism between male and female subtypes specifically in response to injury by 4 weeks. Using a less stringent analysis for sexual dimorphism, we do detect within subtype differences. These are primarily seen in A*β*-RA + A*δ*-LTMRs, and is partly driven by baseline sex differences at a subpopulation-level, which were initially discussed in Fig 3. The variability in low count genes adds noise to these analyses: even deeper sequencing and *in situ* validation may still reveal a clearer interaction, or lack-there-of between sex and injury.

## Discussion

The availability of subtype-specific transgenic mice paired to advances in low input RNA-seq has allowed deep sequencing of sensory neuron subpopulations after injury. This permits detection of a large number of DEGs not currently possible through traditional sc/snRNA-seq approaches. Building on previously available naïve data from Zheng *et al* 2019, we are able to explore sexual dimorphism in naïve states as well as changes after SNI at two timepoints. We have further probed sexual dimorphism after injury at a subpopulation-level.

### Injury Signatures

Samples cluster primarily by cell type. By collapsing these subtypes we can extract general injury signatures that mirror injury signatures seen in previous studies. There is an acute upregulation of key injury genes which decrease over time, along with the more chronic upregulation of genes like *Npy*.

This general analysis is enriched for sensory neurons, as opposed to general whole DRG sequencing. We anticipate this transcriptional signature to be similar to bulk RNA-seq of MACS-purified neurons, where researchers quantified a “nociceptor” transcriptome based on the isolation of small diameter neurons (***Thakur et al. (2014)***). In naïve states, we see many overlapping genes, including key transcription factors they report to be enriched in their “nociceptor” sample, opposed to, whole DRG. These include *Pou4f2*, *Myt1*, *Ldb2*, *Isl2*, *Bhlha9*, and *Atf3*, which show varying degrees of cell-type specificity in our data. We have built on this with added data after injury, giving insight to possible nociceptor-enriched transcription factors involved in injury. For example, *Isl2* encodes Insulin related protein 2 and is regulated after injury in our dataset. It also shows regulation in human patients with diabetic peripheral neuropathy, suggesting cross-species, cross-model target for future experiments (***Hall et al. (2022)***).

To probe differences in acute injury and later states, we examined samples at 3 days and 4 weeks. When comparing all ipsilateral samples, we see a reduction in NP and C-LTMR enrichment more chronically. Population data was pooled for GSEA analyses across timepoints, so changes in gene signatures can be difficult to interpret. For example, if we are working with the assumption that injured NP cells die after SNI, as suggested in West *et al*. 2020, and hinted at by the loss of IB4-binding terminals after nerve injury in rat (***Bailey and Ribeiro-Da-Silva (2006)***), the ipsilateral NP samples by 4 weeks are likely to contain primarily intact neurons (opposed to cell bodies from transected afferents). The significant reduction of an NP signature over time may thus result from changes in the intact NP neurons, or a bias in the general nociceptor population, which shows enrichment for both PEP and NP nociceptors in a naïve state. C-LTMRs show a similar pattern to NP, although we have not queried the loss of this population to the same extent as NP. Together, this is an interested area for future follow-up, as cell loss has also been documented in human patients with neuropathic pain (***Hall et al. (2022)***). To address this in further detail, and amplify a major strength of the study, subtype-specific analyses were performed.

### Subtype-specific injury changes

Previous work has explored injury signatures in whole DRG and single cells. We are adding a middle ground of “bulk", subtype-specific population analyses through deep sequencing of neuronal subtypes post-SNI. Deep sequencing allows us to interrogate genes at a larger dynamic range, including lowly expressed genes, and produces an expression matrix that is less sparse than sc/snRNA-seq.

Our sequencing depth permits differential expression testing within subpopulations (Figure 5A-D). All populations show differential gene expression, but this is primarily driven by general nociceptors and PEP at 3 days, and A*β*-RA + A*δ*-LTMRs at 4 weeks. Using an LFC cutoff of 1, we see significant upregulation of multiple injury genes, and a significant enrichment of a general injury signature in the ipsilateral samples across all subtypes. This suggests the low number of DEGs in other populations is not an artifact of sequencing intact neurons, but instead a biologically relevant signature. A number of these DEGs are involved in gene regulation, and may be useful targets for genetic manipulation.

In line with previous reports, we also see a reduction in cell-type specificity within our injured samples (Figure 5E-I). Primarily, we see a reduced signature in general nociceptors, as well as NP nociceptors, while PEP nociceptors still show enrichment for PEP, either from the contribution of intact afferents in the samples, or less change in injured cells.

There is an added enrichment for A*δ*-LTMRs across all nociceptor subtypes, which remains enriched in LTMR populations. The consequence of this is unclear, with multiple, non-mutually-exclusive hypotheses available. For example, do nociceptors develop a more LTMR-like signature? Is this driven by a clear subset of genes? Is this signature simply representative of a more general, undefined or immature sensory neuron? Do A*δ*-LTMRs show a more injured phenotype in naïve states? We cannot conclusively exclude this latter possibility, but key injury genes are not present in the gene set curated from Zheng and colleagues, and our contralateral *Ntrk2* samples are negatively enriched for our general injury signature at both timepoints. Together, this suggests they do not show a strong injury phenotype at baseline.

### Gene signatures

Using injury signatures extracted from DEG lists we can compare subtype responses over time. By four weeks, *Ntrk2* samples show a distinct injury phenotype not captured by the other populations (Figure 4–5). Loading values suggest a strong involvement of *Car3* and *Car1*, which are both involved in oxidative phosphorylation. Neuropeptide Y (*Npy*) and cerebellin-2 (*Cbln2*) are also highly ranked. These have previously been implicated in mechanical hypersensitivity (***Sandor et al. (2018)***). *Npy* has also been well documented for its upregulation in large-diameter neurons after injury, which may explain the subtype-specific effect seen here (***Wakisaka et al. (1991)***).

### Sexual dimorphism

In a naïve state, we see distinct sexual dimorphism across subtypes (Figure 3). This does not translate to a strong interaction of sex and injury, as the injury response seems to be consistent across sexes (Figure 6). With baseline differences in gene expression across sexes, a strong interaction with injury is not required for functionally relevant changes in injured states. The differences seen in control states may still contribute to painful states due to altered immune/glial interactions, baseline excitability, or differences in higher order circuitry.

Using an additive model to contrast sex and injury at 4 weeks, we are able to identify possible gene candidates for further validation. These represent a set of genes who’s different responses between sexes cannot be solely explained by the general injury response (Figure 6). A small subset shared across populations, which appear to be larger sex-linked, or key injury genes. Of these, *Sprr1a* was previously noted to be sexually dimorphic in naïve states, but also is strongly upregulated after injury, in line with previous reports.

Proportionally, A*β*-RA + A*δ*-LTMRs show the most DEGs when contrasting sex and injury condition, in keeping with a naïve state. One third of these genes were previously reported to differ in control samples, with 12% regulated in injury at 4 weeks. As a top hit, *Slit3* is an estrogen-sensitive axonal guidance molecule previously discussed in the context of endometriosis and pelvic pain (***Greaves et al. (2014)***). Other DEGs include a range of genes involved in inflammation and immune response (eg. *Ifi211*, *Ctla2a*, *Tlr4*), cholinergic receptors (*Chrna3*, *Chrnb3*), the transcription factor *Neurog3*, and numerous sex-linked genes.

### Technical limitations

This data does not conclusively highlight an interaction of sex and injury within subtypes. The absence of a strong signature fits previous literature, but nuanced changes in lowly expressed genes may still hold biological relevance. Tamoxifen dosing introduces a confound so candidates were validated on naïve, wildtype tissue, and the variability in low count genes makes this difficult to probe with our sequencing depth. Shrinkage methods differ in their LFC estimates at this point, with a conservative shrinkage by *apeglm* showing no changes, while a more relaxed shrinkage by *ashr* captures large fold changes, some of which are driven by single samples. To give more confidence to DEGs captured by *ashr*, more stringent filtering and posthoc validation may be warranted to select candidates for external validation.

## Conclusions

Here, we present the deep sequencing of male and female DRG subtypes after SNI as a resource to the field (https://livedataoxford.shinyapps.io/drg-directory/). We show that in addition to stereotyped changes after injury, neuronal populations undergo subpopulation-specific changes at a molecular level, and these vary with time. In naïve states, we see subpopulation-specific sexual dimorphism that is retained in injured states. Taken together, this data provides a starting point for future experimentation surrounding subpopulation differences as well as stereotyped changes across timepoints, and highlights the importance of factoring sex into these studies.

## Methods and Materials

All work was done in accordance with the UK Home Office and the University of Oxford Policy on the Use of Animals in Scientific Research. This study conforms to ARRIVE guidelines.

Animals were housed in standard conditions on a 12-hour light/dark cycle with food and water *ad libitum*. All animals were randomly assigned to experimental groups where applicable. Internal controls were used when not possible to randomize (ie. ipsilateral vs contralateral comparisons). Unless explicitly stated, all experiments were performed on both males and females. Briefly, driver lines were bred with a fluorescence reporter for various experiments. When necessary, inducible lines were dosed with intraperitoneal (i.p.) injection(s) of tamoxifen. Specific details below.

### Transgenic details

C57BL/6 mice were purchased from the Oxford University Breeding Unit. Cre driver lines used include: Calca^tm1.1(cre/ERT2)Ptch^ (CGRP, gifted from Prof. Pao-Tien Chaung) ***Song et al. (2012)***, Mrgprd^tm1.1(cre/ERT2)Wql/J^ (MRGPRD, JAX 031286) ***Olson et al. (2017)***, Scn10a^tm2(cre)Jnw^ (Nav1.8, gifted from Prof. John Wood) ***Nassar et al. (2004)***, Th ^tm1.1(cre/ERT2)Ddg/J^ (TH, gifted from Prof. David Ginty) ***Watanabe et al. (2017)***, Ntrk2^tm1.1(cre/ERT2)Ddg/J^ (TRKB, gifted from Prof. Paul Heppenstall) ***Dhandapani et al. (2018)***. Cre-driver lines were bred and maintained as heterozygotes, except for Th^creERT2^, which was bred as homozygous. Details are listed in Table 1.

The following reporters were used to visualize DRG subpopulations: B6.129S-Gt(ROSA)26Sor^tm32(CAG-COP4*H134R/EYFP)Hze/J^ (JAX 012569, gifted from Prof. Simon Butt), tdTomato B6.Cg-Gt(ROSA)26Sor^tm14(CAG-tdTomato)Hze/J^ (JAX 007914) and ai80D B6.Cg-Gt(ROSA)26Sor^tm80.1(CAG-COP4*L132C/EYFP)Hze/J^, (JAX 025109). Ai32 and ai14 use was based on initial breeding availability. Ai80 depends on both Flp- and Cre-recombinase for intersectional targeting, and was used for neuronal targeting of *Ntrk2*. Ai80 was first crossed to an Advillin^flpO^ (unpublished, gifted from Prof. David Ginty). Reporters were bred as homozygotes where applicable. Advillin^flpO^ was bred to a C57BL/6 background for at least 7 generations prior to experimental use.

### Tamoxifen regimes

Tamoxifen (Sigma-Aldrich) was dissolved 20 mg/ml in corn oil via sonification. All animals were dosed i.p. and health statuses were monitored daily for the duration of the dosing regime. Calca^creERT2^ were dosed 5x (daily) with 75 mg/kg in adulthood. Mrgprd^creERT2^ were dosed 5x i.p. (0.5 mg/animal/day), beginning between P10-P17. Body weight recovered more quickly when dosed at later stages, with no noticeable difference in reporter expression. We recommend dosing begin at P17 for this line moving forward. Th^creERT2^ were dosed 1x with 50 mg/kg above 6 weeks of age. Ntrk2^creERT2^ were dosed 5x (daily) with 75 mg/kg in adulthood.

### Spared nerve injury

Adult mice were anesthetized with 2% inhaled isoflurane. Using sterile technique (including incision site sterilization and surgical drapes), the sciatic nerve was exposed prior to ligation and transection of the tibial and common peroneal branches ***Decosterd and Woolf (2000)***. The sural nerve was left intact. Each animal was dosed with systemic (5 mg/kg Rimadyl, Pfizer) and local (2 mg/kg Marcain, AstraZeneca) postoperative analgesia. Animals were monitored daily for self-mutilation, and no animals required sacrifice due to tissue damage.

### Sample collection

Sample size was calculated using the algorithm published by Zhao et al. (***Zhao et al. (2018)***). See supplemental methods for full details. In total, 160 paired samples were collected (ipsilateral and contralateral) over five neuronal subtypes at an acute (3 day) and late (4 week) timepoint after SNI. Ipsilateral lumbar (L3-L5) were compared to contralateral (L3-L5) DRGs to ensure an internal control. One animal was used for each sample pair, excluding Th^creERT2^, where DRGs from two animals were pooled during dissection. Male and female samples were evenly split, with the exclusion of Mrgprd^creERT2^, 3 days after SNI where only 3 female mice could be used. Five males were thus processed for this group.

Multiple animals were processed in parallel but collection times from perfusion to frozen were kept to less than 4 hours. Adult animals were first overdosed with pentobarbital and perfused transcardially with sterile, ice cold saline. Lumbar DRG were quickly removed and placed into HBSS on ice. Post-dissection of all tissue, collagenase/dispase was added for a 60 min digest at 37 C followed by mechanical dissociation with polished glass pipettes. Myelin and debris was reduced using a clean 15% w/v BSA cushion. Samples were placed on ice and centrifuged at 4 C as much as possible (i.e. excluding digestion). Prior to FACS, a subset of neurons from each sample was examined under brightfield.

### Library preparation and sequencing

Samples were transferred on ice immediately to the WIMM FACS Facility (Oxford) for sorting on a BD FACSAria Fusion 1 or Fusion 2. For each condition, 100 cells were isolated directly into low protein binding eppendorfs containing 2 ul NEBNext Single Cell Lysis Buffer (NEB, E5530S). Samples were kept on dry ice until transfer to −80 C for overnight storage.

Once all samples were collected, samples were thawed on ice, vortexed, and randomized into a 384-well 4titude Framestar skirted PCR plate (Brooks Life Science, 4ti-0384/C; Thermo Scientific, AB-0558). Non-directional libraries were prepared together using NEB Ultra low/Smarter library prep, as per manufacture instructions by the Oxford Genomics Centre at the Wellcome Trust Centre for Human Genetics. Libraries were amplified (21 cycles) on a Tetrad (Bio-Rad) using in-house unique dual indexing primers (based on ***Lamble et al. (2013)***). Individual libraries were normalized using Qubit, and the size profile was analysed on the 2200 or 4200 TapeStation before pooling together accordingly. The pooled library was diluted to 10 nM for storage. The 10 nM library was denatured and further diluted prior to loading on the sequencer. Sequencing was performed over three independent runs, and merged after quality control. Paired end sequencing was performed using a NovaSeq6000 platform using the S2/S4 reagent kit v1.5. Samples were sequenced with a 150 bp read length, at a depth of 30 million reads per sample. Raw data is available on GEO (GSE216444).

### Analysis

#### Overview

Reads were mapped to the GRCm38 (mm10) Mouse Genome using STAR alignment (***Dobin and Gingeras (2013)***). Samtools was used to sort, index, and merge BAM files (***Li et al. (2009)***). Quality control (QC) was performed with both FastQC and Samtools prior to gene counting with HTSeq (***Andrews (2010)***; ***Li et al. (2009)***; ***Anders et al. (2015)***). Software is listed in ***Table 2***.

#### Quality control

Samples were judged based on library size, as well as read assignment, alignment, and normalized gene coverage. Together, six samples were removed from downstream analyses, with details in the supplemental.

#### DESeq2

Counts were corrected for effective library size in R using DESeq2 (***Love et al. (2014)***). Normalized gene counts were fitted to a negative binomial distribution. A batch effect was introduced during sample collection, and a model that included this batch effect was fitted to every gene. The significance of the model’s coefficients was assessed using the Wald test.

Counts were log transformed via variance stabilizing transformation (VST). VST transformed counts were used for all plotting, unless otherwise stated. The R package Limma was used to remove the batch effect in PCA and heatmap figures. Uncorrected PCA plots are shown in Supplemental Figure 2. Box plots show median + interquartile range (IQR), with 1.5*IQR whiskers. Principal component analysis (PCA) was performed using the top 5000 ENSEMBL genes ranked by standard deviation. Sample distances are proportional to Mahalanobis distance, and ellipses show the 95% confidence interval of a condition’s gene expression distribution. Hierarchical clustering was done on transformed counts using Euclidean distances and complete linkage. Gene enrichment within neuronal subtypes was calculated using VST counts, and was defined as genes with a subpopulation mean within the top 75% of expressed genes, across contralateral samples.

#### Custom GSEA

Deep RNA-seq of naïve DRG subpopulations has been previously performed elsewhere (***Zheng et al. (2019)***). These results were curated into subpopulation-enriched gene sets to probe enrichment in the current data, with full details in the supplementals. Briefly, RNA-seq count data was accessed from https://www.ncbi.nlm.nih.gov/geo/ (GSE131230). Expression data were generated via STAR alignment and HTSeq on the same genome build. Counts were corrected for library size and transformed via rlog in R using DESeq2 and filtered to match their published report. Within each subpopulation, genes with a mean rlog above the 95% quantile cut-off were curated into a ‘gene set’ for subpopulation enrichment. These custom gene sets were then compiled for a GSEA analysis to our contralateral (“naïve"-like) samples using the clusterProfiler package in R using ranked fold changes from mean(subpopulation)/mean(total).

#### Differential expression testing

Differential expression testing was performed on filtered data using the Wald test and a weighted FDR correction (independent hypothesis weighting, IHW). Effect sizes were calculated using Bayesian shrinkage estimators (the *apeglm* method, via DESeq2) and are presented as moderated (shrunken) log_2_ fold changes (LFC) (***Zhu et al. (2019)***), with full details in the supplementals. Significance was set at an FDR < 0.05 and a LFC > 1.

GSEA analyses against “all gene sets” were performed using ranked log_2_ fold changes (LFC) via msigdbr (***Dolgalev (2021)***) and clusterProfiler (***Wu et al. (2021)***) libraries. Custom GSEA analyses were calculated against a curated list of enriched genes from previously published subpopulation data, as described (***Zheng et al. (2019)***). Ipsilateral sample enrichment was calculated against the same combined contralateral baseline above. GO term analyses for differentially expressed genes (DEGs) were performed using the Wallenius method via goSeq (R) (***Young et al. (2010)***). The filtered count data of expressed, non-DEG genes were used as a background. Protein interaction networks were generated using STRING (***Szklarczyk et al.(2021)***).

#### Injury signature enrichment

Acute and late injury signatures were calculated using supervised principal component analyses (SPCAs) ***Bair et al. (2012)*** on DEGs at 3 days or 4 weeks from the general injury analysis. Eigenvectors were extracted from the first principal component (PC1) and correlated across samples as an unbiased injury signature. For the 4 week timepoint, PC2 was also analysed, and loading values were also extracted. These are a product of the covariance between the scaled dimensions and the original variables, giving a weight to how much individual genes contribute to each principal component.

### Data accessibility

This dataset highlights molecular changes in sensory neuron subtypes across multiple timepoints in a murine neuropathic pain model. To improve the accessibility of this data, an open-source database is available at https://livedataoxford.shinyapps.io/drg-directory/, which includes shared code to generate personal -omics hosting sites. Raw and processed count data are available on GEO, reference GSE216444.

### Tissue staining (IHC and *in situ*)

Adult animals were overdosed with pentobarbital and perfused transcardially with sterile saline followed by 4% paraformaldehyde. Tissue for immunohistochemistry was removed and post-fixed prior to subsequent dehydration in 30% sucrose (0.1M PB) at 4 C for a minimum of 48 hours. Samples were then embedded in OCT medium (Tissue-Tek), sectioned, and stored at −80 C. Neuronal profiles were quantified across multiple sections per animal, opposed to more detailed stereology, and are presented as estimates. In addition to subpopulation markers such as TH (C-LTMRs), CGRP (peptidergic) and parvalbumin (PV, proprioceptors), non-peptidergic neurons bind isolectin B4 (IB4) from Griffonia simplicifolia. Neurofilament heavy chain (NF200) labels large diameter neurons in mice, and NeuN (or FOX3) is a general neuronal marker.

To validate sex differences in the absence of tamoxifen dosing, fresh DRG were isolated from littermate pairs of male and female wildtype mice (n=3, multiple sections per mouse). Tissue was embedded directly in OCT prior to freezing on dry ice and storage at −80 C. *In situ* hybridizations (ISH) were performed using RNAscope Multiplexing v1 and v2 as per manufacturer’s instructions (ACDBio) and TSA Vivid fluorophores (7526/1, 7523/1), with probe details in Table 3.

## Supporting information

Supplemental Table 1. Gene enrichment by population.

Supplemental Table 2. GSEA enrichment scores for naive validation.

Supplemental Table 3. Gene Sets based on Zheng et al 2019

Supplemental Table 4. DEG naive sex

Supplemental Table 5. DEG overlap naive sex

Supplemental Table 6. GO naive sex

Supplemental Table 7. GSEA naive sex

Supplemental Table 8. DEG general injury

Supplemental Table 9. DEG overlap general injury

Supplemental Table 10. GO general injury

Supplemental Table 11. GSEA general injury

Supplemental Table 12. GSEA enrichment scores for general injury

Supplemental Table 13. DEG subtype injury

Supplemental Table 14. DEG subtype injury, opposing direction

Supplemental Table 15. GO subtype injury

Supplemental Table 16. DEG subtype injury. Gene regulation DEGs

Supplemental Table 17. DEG subtype injury. Transcription factor DEGs

Supplemental Table 18. GSEA enrichment scores for subtype injury

Supplemental Table 19. DEG additive model sexual dimorphism

## Acknowledgments

This work was funded in part by the Wellcome Trust (DPhil scholarship to AMB, 215145/Z/18/Z) and a Wellcome Investigator Grant to DB (223149/Z/21/Z), as well as the MRC (MR/T020113/1), and with funding from the MRC and Versus Arthritis to the PAINSTORM consortium as part of the Advanced Pain Discovery Platform (MR/W002388/1). AMB further received a GTC MSDTC Scholarship. GB is funded by Diabetes UK, grant number 19/0005984, MRC and Versus Arthritis through the PAINSTORM consortium as part of the Advanced Pain Discovery Platform (MR/W002388/1) and by the Wellcome Trust (223149/Z/21/Z). XY was partly funded by Shanghai Key Laboratory of Peripheral Nerve and Microsurgery (20DZ2270200), NHC Key Laboratory of Hand Reconstruction (Fudan University), Shanghai, China.

This research was funded in part by the Wellcome Trust [215145/Z/18/Z and 223149/Z/21/Z]. For the purpose of open access, the author has applied a CC BY public copyright license to any Author Accepted Manuscript version arising from this submission.

## Supplemental Methods

### Sample collection and mapping

Sample size was calculated using the algorithm published by Zhao *et al*. (***Zhao et al. (2018)***). Based on our sequencing depth, we expect to detect 10000 genes per sample. Recent data from Parisien and colleagues using whole DRG suggests that 1000 genes can be considered regulated (based on an FDR cutoff of 0.05) in DRG one week after SNI, with an average fold change of 3.3 (SNI/control) (***Parisien et al. (2019)***). Prior data collection in our lab indicates that the minimum average read counts among the prognostic genes is 700 and the maximum expected dispersion is 0.2. Using a desired minimum fold change set to 3.3, we can achieve a power of 0.99 with 8 samples per condition (minimum read count = 700, dispersion = 0.2, average fold change = 3.3, FDR = 0.05). Moreover, we will be sufficiently powered to detect large fold changes between sexes (power of 0.8, n = 4). All surgeries and sample collections were performed by the same experimenter to reduce variability across samples.

Reads were mapped to the GRCm38 (mm10) Mouse Genome using STAR alignment with the parameters below (7) (***Dobin and Gingeras*** (***2013***)). Samtools was used to sort, index, and merge BAM files ***Li et al. (2009)***. Quality control (QC) was performed with both FastQC and Samtools prior to gene counting with HTSeq (***Andrews (2010)***; ***Li et al. (2009)***; ***Anders et al. (2015)***).

**Appendix 0—figure 7.**
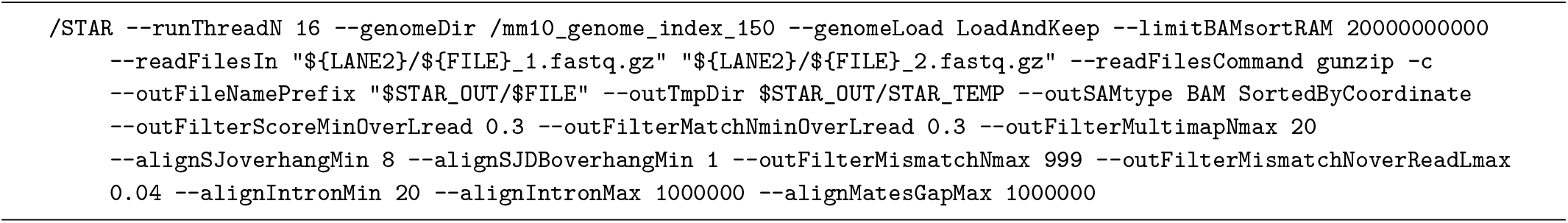
STAR alignment

### Quality control

Samples were judged based on raw count numbers, as well as read assignment, alignment, and normalized gene coverage. Together, six samples were removed from downstream analyses. Two samples immediately failed to sequence, as clearly highlighted by the lack of reads. Primary alignments to the genome showed consistency across samples, and STAR mapping by sample per sequencing run shows consistency across sequencing run.

Normalised position against coverage was used to measure 3’ prime bias, as this is often the result of RNA degradation. Here, a few samples show reduced coverage at the 5’ end. Samples which showed bias coverage towards the 3’ end and had lower genome alignment to coding regions were removed from downstream analyses. High-level QC was then performed through cluster analyses and gene mapping using DESeq2 (below).

### DESeq2

Counts were corrected for effective library size in R using DESeq2 (***Love et al. (2014)***). Normalized gene counts were fitted to a negative binomial distribution. A batch effect was introduced during sample collection, as discussed below. A model that included this batch effect was fitted to every gene and the significance of the model’s coefficients was assessed using the Wald test.

Counts were transformed via variance stabilizing transformation (VST). VST transformed counts were used for all plotting, unless otherwise stated. The R package Limma was used to model the batch effect in PCA and heatmap figures. Uncorrected plots are shown in the chapter appendix. Box plots show median + interquartile range (IQR), with 1.5*IQR whiskers. Principal component analysis (PCA) was performed using the top 5000 ENSEMBL genes ranked by standard deviation. Sample distances are proportional to Mahalanobis distance, and ellipses show the 95% CI of a conditions gene expression distribution. Hierarchical clustering was done on transformed counts using Euclidean distances and complete linkage. Gene expression dot plots were created using median VST counts, and coloured by gene expression. Dot sizes are calculated as exponential VST counts to reflect differences in more highly expressed genes. Due to the range of base means in sexually dimorphic genes, dot sizes here match VST counts. Gene enrichment within neuronal subtypes was calculated using VST counts, and was defined as genes with a subpopulation mean within the top 75% of expressed genes, across contralateral samples.

Heatmaps were generated from transformed counts and visualized using “complex heatmaps” (***Gu et al. (2016)***). GSEA analyses were performed using ranked log_2_ fold changes via msigdbr (***Dolgalev (2021)***) and clusterProfiler (***Wu et al. (2021)***) libraries.

### Custom GSEA

Deep RNA-seq of naïve DRG subpopulations has been previously performed elsewhere ***Zheng et al. (2019)***. Eight DRG subpopulations were studied after transgenic labelling and fluorescent sorting, with 3-5 samples per population (mixed sex, multiple mice per sample). These populations include non-peptidergic (NP) and peptidergic (PEP) nociceptors, C-LTMRs, A*δ*- LTMRs, A*β*-SA1-LTMRs, A*β*-Field-LTMRs, A*β*-RA-LTMRs, and proprioceptors (PROP). Full methodology can be found in their published report (***Zheng et al. (2019)***).

These results were curated into subpopulation-enriched gene sets to probe enrichment in the current data. RNA-seq count data was accessed from https://www.ncbi.nlm.nih.gov/geo/ (GSE131230). Expression data was generated via STAR alignment and HTSeq on the same genome build, mirroring the current study ***Zheng et al. (2019)***. Counts from Zheng and colleagues were then corrected for library size and transformed via rlog in R using DESeq2 to match their published report. Principal component analysis (PCA) was performed using the top 5000 ENSEMBL genes ranked by standard deviation. Genes were then filtered across all samples for a mean rlog greater than 0, to mirror their filtering criteria. From this filtered dataset, 95% quantiles were determined for each gene. Within each subpopulation, genes with a mean rlog above the 95% quantile cut-off were curated into a ‘gene set’ for subpopulation enrichment. These custom gene sets were compiled for a GSEA analysis to our contralateral (“naïve"-like) samples using the clusterProfiler package in R. Here, GSEA analyses were performed using ranked fold changes from mean(subpopulation)/mean(total) VST counts per contralateral samples.

### Within population sex differences

Recent spatial-seq data has highlighted within-population sex differences in humans (***Tavares-Ferreira et al. (2022)***). The current study is well powered to explore this in murine DRG populations.

Hypothesis testing for each population was performed on genes with at least 10 combined reads across all contralateral samples (35609 ENSEMBL genes) and was performed using the Wald test and a weighted FDR correction (independent hypothesis weighting, IHW). IHW first weights p-values by mean expression, before correcting for multiple hypothesis testing by Benjamini Hochberg (BH). This increases our power to detect differences in highly expressed genes. A Cook’s distance cutoff was set as the .99 quantile to filter genes with excessive variation. Effect sizes were calculated using Bayesian shrinkage estimators (the *apeglm* method, via DESeq2) and are presented as moderated (shrunken) log_2_ fold changes (LFC) (***Zhu et al. (2019)***).

Sixteen samples were processed across time points for each population, with 8 male and 8 female samples each. The NP sample set is composed of 7F + 9M, due to breeding availability. Only 15 samples passed QC for nociceptors (7M + 8F).

Gene ontology (GO) was performed on differentially expressed genes in R using the goSeq library (FDR < 0.05, LFC > 1) (***Young et al. (2010)***). P-values were calculated by the default Wallenius method. Here, filtered count data of non-DEG genes were used as a background, with a cut-off requiring 10 reads across samples. clusterProfiler (R) was used for GSEA analyses, with BH correction.

### Injury modelling

Two design models were fitted and used to examine injury changes - with and without subtype information. In each case, batch was included. Design models were compared against reduced models using the likelihood ratio test (LRT), and the distribution of variance per gene by factor was calculated using the package variancePartition (***Hoffman and Schadt (2016)***).

### Differential expression analyses

Differential expression testing was performed on filtered data using the Wald test and a weighted FDR correction (independent hypothesis weighting, IHW). Count data was first filtered to genes with an average of 5 reads in at least 10% of the samples.

GSEA analyses against “all gene sets” were performed using ranked log_2_ fold changes (LFC) via msigdbr (***Dolgalev*** (***2021***)) and clusterProfiler (***Wu et al. (2021)***) libraries. Custom GSEA analyses were calculated against a curated list of enriched genes from previously published subpopulation data Updating. Ipsilateral sample enrichment was calculated against the same combined contralateral baseline. GO term analyses for DEGs were performed using the Wallenius method via goSeq (R). The filtered count data of expressed, non-DEG genes were used as a background (here, 5 reads in at least 10% of the samples). Protein interaction networks were generated using STRING (***Szklarczyk et al. (2021)***).

### Injury phenotypes

To compare the general effect of SNI across early and later states, condition (ipsi vs cont) and time (3D vs 4W) were modelled as a grouping factor in an additive design, combining samples across neuronal subtypes. Testing within subtypes was performed using a grouping factor for time, condition, and population.

### Sexual dimorphism

Sexual dimorphism in injured samples was interrogated within populations using an interaction model, as well as by contrasting sex and injury from an additive design. Technically, the second method requires a shift to a different Bayesian shrinkage estimator, from “approximate posterior estimation for the general linear model” (*apeglm*) using a heavy-tailed Cauchy prior distribution to a more generic adaptive shrinkage method (*ashr*) (***Love et al. (2014)***). While both are widely accepted shrinkage methods, the consequences of this shift are discussed throughout the results.

- Supplemental Table 1. Gene enrichment by population.
- Supplemental Table 2. GSEA enrichment scores for naïve validation.
- Supplemental Table 3. Gene Sets based on Zheng *et al*. 2019.
- Supplemental Table 4. DEG naïve sex.
- Supplemental Table 5. DEG overlap naïve sex.
- Supplemental Table 6. GO naïve sex.
- Supplemental Table 7. GSEA naïve sex.
- Supplemental Table 8. DEG general injury.
- Supplemental Table 9. DEG overlap general injury.
- Supplemental Table 10. GO general injury.
- Supplemental Table 11. GSEA general injury.
- Supplemental Table 12. GSEA enrichment scores for general injury.
- Supplemental Table 13. DEG subtype injury.
- Supplemental Table 14. DEG subtype injury, opposing direction.
- Supplemental Table 15. GO subtype injury.
- Supplemental Table 16. DEG subtype injury. Gene regulation DEGs.
- Supplemental Table 17. DEG subtype injury. Transcription factor DEGs.
- Supplemental Table 18. GSEA enrichment scores for subtype injury.
- Supplemental Table 19. DEG additive model sexual dimorphism.

---

**Appendix 0 —table 1.**
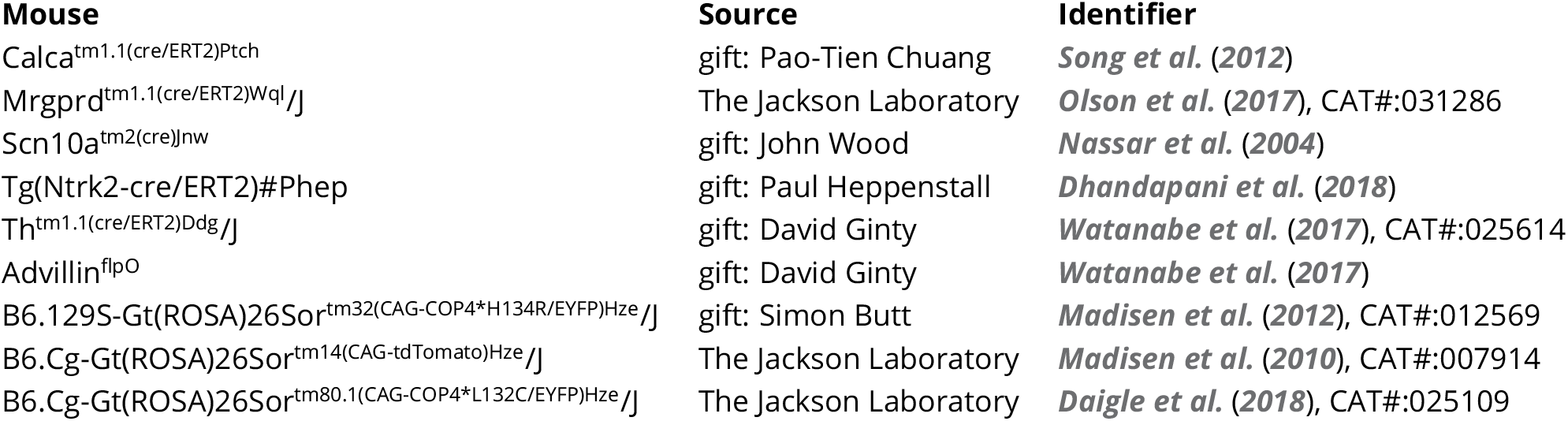
Transgenic lines used in the current study

**Appendix 0 —table 2.**
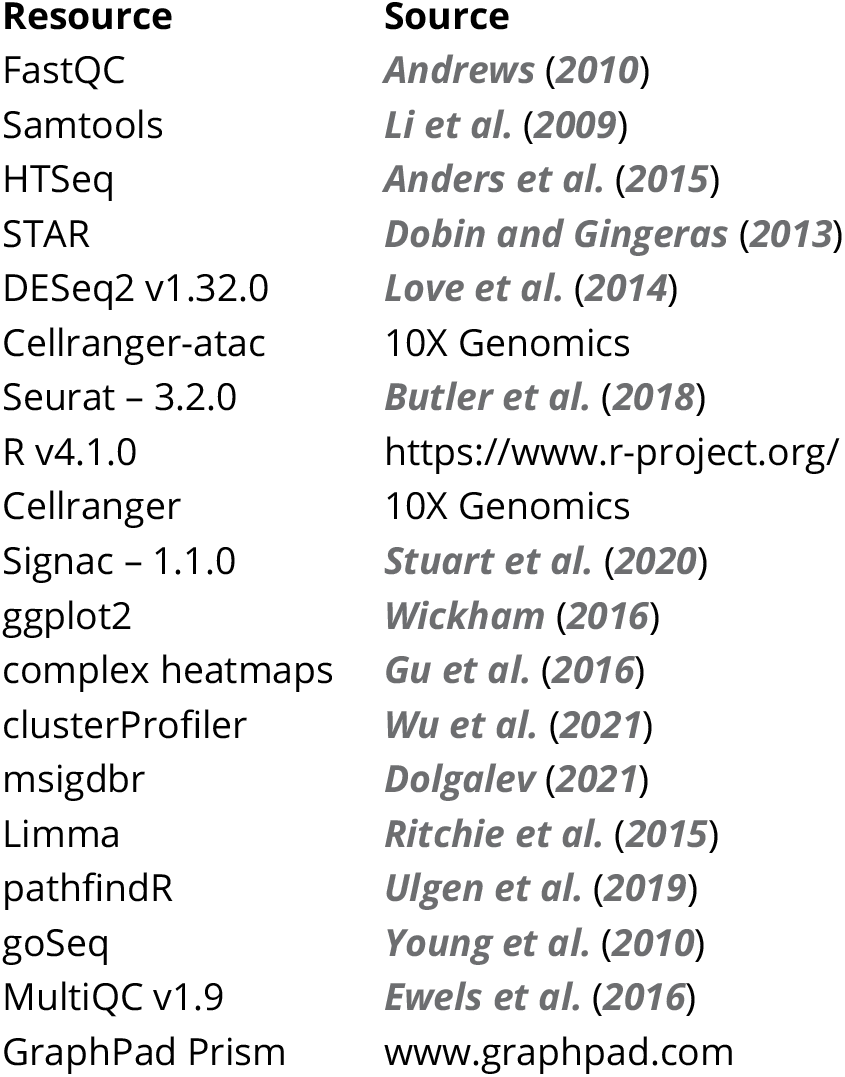
Software used in the current study

**Appendix 0 —table 3.** Commercial reagents used in the current study

| Reagent | Source | Identifier | Dilution |
| --- | --- | --- | --- |
| Donkey anti-Mouse IgG (H+L), Alexa Fluor 488 | ThermoFisher Scientific | A-21202 | 1/200 |
| Donkey anti-Rabbit IgG (H+L), Alexa Fluor 546 | ThermoFisher Scientific | A10040 | 1/200 |
| Donkey anti-Sheep IgG (H+L), Alexa Fluor 488 | ThermoFisher Scientific | A-11015 | 1/200 |
| Donkey anti-Sheep IgG (H+L), Alexa Fluor 546 | ThermoFisher Scientific | A-21098 | 1/200 |
| Goat anti-Chicken IgY (H+L), Alexa Fluor 546 | ThermoFisher Scientific | A-11040 | 1/200 |
| Goat anti-Rabbit IgG (H+L), Alexa Fluor 488 | ThermoFisher Scientific | A-11008 | 1/200 |
| Goat anti-Rabbit IgG (H+L), Alexa Fluor 546 | ThermoFisher Scientific | A-11010 | 1/200 |
| Goat anti-Rabbit IgG (H+L), Pacific Blue | ThermoFisher Scientific | P-10994 | 1/200 |
| Streptavidin, Pacific Blue conjugate | ThermoFisher Scientific | S11222 | 1/100 |
| anti-CGRP, rabbit | Peninsular Labs | T4032 | 1/500 |
| anti-CGRP, sheep | Enzo | Ca1137 | 1/250-1/500 |
| anti-Glutamine synthetase (GS), rabbit | Sigma-Aldrich | G2781 | 1/500 |
| anti-NeuN, chicken | Merck Millipore | Abn91 | 1/500 |
| anti-NeuN, rabbit | Abcam | Ab177487 | 1/500 |
| anti-NF200, mouse | Sigma-Aldrich | N0142 | 1/250 |
| anti-NF200, rabbit | Merck Millipore | ABN76 | 1/1000 |
| anti-Parvalbumin (PV), guinea pig | Frontier Institute | Af1000 | 1/200-1/500 |
| anti-Tyrosine Hydroxylase (TH), sheep | Merck Millipore | Ab1542 | 1/400 |
| IB4, streptavidin conjugated | Sigma-Aldrich | L2140 | 1/100 |
| mm-Ntrk2-C2 | ACDBio | 423611-C2 | 1/50 |
| mm-Kcnj11 | ACDBio | 431451 | NA |
| mm-Kcns1 | ACDBio | 525941 | NA |
| mm-Cacng2 | ACDBio | 437221 | NA |
| RNAscope® Multiplex Fluorescent Reagent Kit v1 | ACDBio | 320293 | NA |
| RNAscope® Multiplex Fluorescent Reagent Kit v2 | ACDBio | 323100 | NA |
| TSA Vivid Fluorophore kit 570 | Biotechne | 7526/1 | 1/1500 |
| TSA Vivid Fluorophore kit 520 | Biotechne | 7523/1 | 1/1500 |

**Figure 1 — figure supplement 1.**
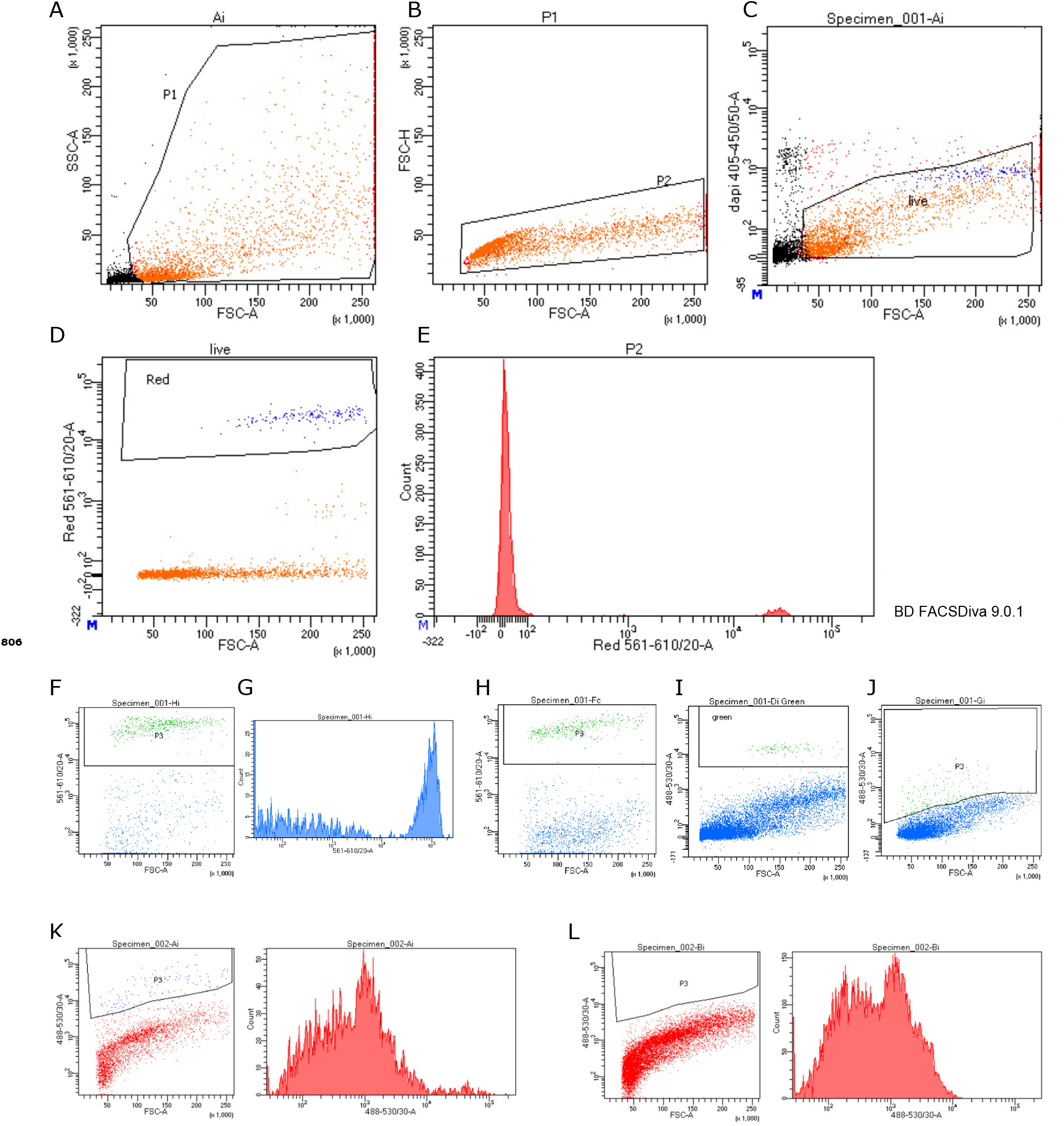
Sensory neuron populations sorted by fluorescence. A-E: Mrgprd^creERT2^ sorted neurons, gated for scatter (A), doublets (B), live/dead (C), and ai14 fluorescence (D) and associated histogram (E). F: Scn10a^cre^ gating for ai14, and associated histogram (G). H: Calca^creERT2^ gating for ai14. I: Th^creERT2^ gating for ai32 reporter. J: Ntrk2^creERT2^ gating for ai80 reporter. K-L: FACS validation for Ntrk2^creERT2^, positive (K) and negative control (L) gated for CatCH/EYFP fluorescence in 488.

**Figure 1 — figure supplement 2.**
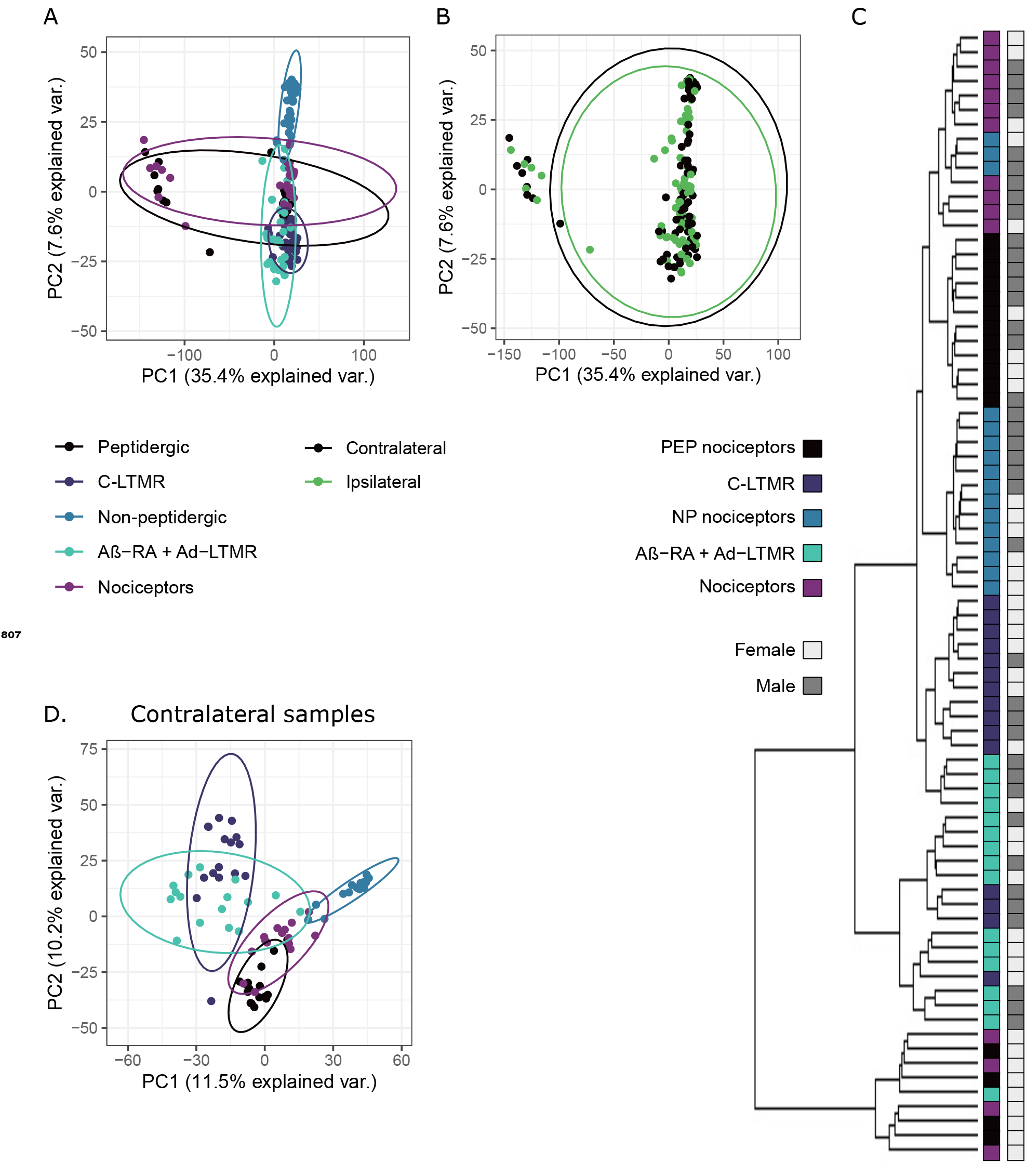
Supplemental sample clustering. A-B. PCA biplot without batch correction by subtype (A) and injury status (B). C. Hierarchical clustering of contralateral samples, annotated by subpopulation and sex. D. PCA-bi-plot of contralateral samples.

